# Differential Burst dynamics of Slow and Fast gamma rhythms in Macaque primary visual cortex

**DOI:** 10.1101/2025.10.01.679813

**Authors:** Vignesh Raju, Supratim Ray

**Affiliations:** Undergraduate Program, Indian Institute of Science, Bengaluru, India, 560012; Centre for Neuroscience, Indian Institute of Science, Bengaluru, India, 560012

**Keywords:** Gamma, oscillations, bursts, LFP, Matching Pursuit

## Abstract

Gamma oscillations have been ubiquitously observed across a wide spectrum of brain areas in multiple species. They tend to occur intermittently in the form of bursts, rather than being produced as sustained and continuous rhythmic activity. Recent studies have shown that large visual sinusoidal gratings elicit two distinct gamma rhythms, namely, slow (≈ 20-35 Hz) and fast gamma (≈ 40-65 Hz), in the primary visual cortex (V1) of non-human primates. However, their mechanisms of generation and potential functional role in cortical processing remain unclear. Details of their burst signatures could potentially provide crucial insights about how the two rhythms influence network dynamics. Therefore, we computed burst statistics (durations and latencies) of simultaneously induced slow and fast gamma rhythms in the local field potential (LFP) recorded from area V1 of two adult female bonnet monkeys using several burst estimation methods. We found that slow gamma rhythm exhibited significantly longer burst durations and longer latencies as compared to fast gamma. Slow gamma exhibited higher long-range synchrony compared to fast gamma, as estimated by coherence and weighted phase lag index (WPLI), which could aid in enhanced global coordination in neocortex. Interestingly, longer burst length of slow-gamma could be replicated in a recently-developed noisy Wilson-Cowan network model by simply changing the firing-rate time-constant of the corresponding inhibitory interneuronal population, which leads to both slower and longer bursts. These results are consistent with the hypothesis that the two oscillations are generated by different inter-neuronal classes that operate over different temporal and spatial scales of integration.

**Significance statement:** Slow (≈ 20-35 Hz) and fast gamma (≈ 40-65 Hz) are two distinct rhythms known to be induced by large visual gratings in the primary visual cortex (V1). Interestingly, gamma oscillations manifest in the form of transient and stochastic epochs, generally termed as “bursts”. We estimated the durations of stimulus-induced slow and fast gamma bursts generated in the local field potential (LFP) recorded from V1 of monkeys. Slow gamma bursts had significantly longer durations and increased latency to onset compared to fast gamma, which was replicated in a noisy Wilson-Cowan model by changing the time-constant associated with inhibitory neuronal population. These results suggest that slow and fast gamma are generated by different inter-neuronal networks operating at different spatio-temporal scales.

## Introduction

Gamma rhythm (typically, 30-70 Hz) has been associated with diverse cognitive phenomena, including memory (Lisman and Idiart, 1995), selective attention (Jensen et al., 2007) and feature binding through synchronisation (Han et al., 2022). Cognitive impairments such as Alzheimer’s Disease (AD), schizophrenia, and Attention-Deficit Hyperactivity Disorder (ADHD) have been correlated with atypical gamma oscillations (Herrmann and Demiralp, 2005; Uhlhaas and Singer, 2010; Murty et al., 2021). Notably, a decline in the power of gamma oscillations, even with normal ageing in healthy individuals, has been recently reported (Murty et al., 2020). They are widely believed to be generated due to network interactions between inhibitory and excitatory populations of neurons.

Experimental studies have revealed the presence of multiple gamma rhythms, potentially mediating distinct information processing pathways, in the mouse hippocampus (Colgin et al., 2009; Belluscio et al., 2012). In non-human primates, large visual gratings have been shown to induce two distinct gamma rhythms: slow (≈ 20-35 Hz) and fast gamma (≈ 40-65 Hz), in the primary visual cortex (V1) (Murty et al., 2018). Recent studies using optogenetics in mouse V1 have elucidated the role of distinct inhibitory neuronal populations, namely somatostatin (SOM) and parvalbumin-positive (PV+) interneurons, in the generation of slow and fast gamma, respectively (Chen et al., 2017; Veit et al., 2017). Furthermore, these distinct gamma rhythms have been linked to selective representation of stimulus features such as spatial frequency and contrast in the striate cortex of rodents (Han et al., 2021b). Slow gamma has been proposed to play a long-range modulatory role, while fast gamma has been implicated to mediate local processing based on their unique phase-amplitude coupling signatures and coherence measurements in monkey V1 (Murty et al., 2018; Prabhu and Ray, 2024).

A separate line of investigation has revealed gamma oscillations to be generated stochastically, in the form of bursts in the macaque V1 (Xing et al., 2012; Chandran Ks et al., 2018). Such brief bursts are observed in multiple bands (such as beta and gamma) and in multiple brain regions, and have been hypothesized to play a role in encoding and retrieval during the working memory paradigm in the prefrontal cortex of monkeys (Lundqvist et al., 2016) and in facilitating information flow across cortical areas on a cycle-by-cycle basis (Palmigiano et al., 2017). Such bursty oscillations have been modelled using Wilson-Cowan (WC) models that are driven by Poisson inputs (Jia et al., 2013) or a non-linear sustained limit cycle-model (Jadi and Sejnowski, 2014) with Ornstein-Uhlenbeck (OU) noise inputs (Krishnakumaran and Ray, 2023). Further, recent models have shown that the burst profiles could be critically influenced by the underlying network connectivity parameters and the appropriate regime of oscillations (Powanwe and Longtin, 2019). Hence, a detailed characterization of burst statistics could convey important information about the underlying neural circuitry.

Here, we aim to characterize the burst statistics, such as duration and temporal distribution, of simultaneously induced slow and fast gamma bursts. We recorded Local Field Potential (LFP) using microelectrodes from the area V1 of two awake adult female bonnet monkeys performing a passive fixation task, while they viewed large-sized sinusoidal gratings that induced both the gamma rhythms. We estimated burst lengths of slow and fast gamma rhythms present in the dataset using the Matching Pursuit (MP) algorithm, an iterative and greedy signal decomposition technique that has been shown to robustly compute burst durations (Chandran Ks et al., 2018). We also used other methods, including traditional methods such as Hilbert Transform as well as a recently developed compressed sensing technique that can detect bursts using smaller but dynamic dictionaries (Anand et al., 2025). Further, we studied the extent of synchronization in slow and fast gamma by analyzing phase coherence and weighted-phase lag index (WPLI) profiles. Finally, we tested whether the burst characteristics could be explained in the OU noise model (Krishnakumaran and Ray, 2023) by varying the firing rate time constant of the inhibitory neuronal population.

## Materials and Methods

### Animal preparation and recording

Two adult female bonnet monkeys (*Macaca radiata*) weighing 4 kg and 3.3 kg were used in this study. Experimental protocols used for the animals were in accordance with the guidelines approved by the Committee for the Control and Supervision of Experiments on Animals (CCSEA) and the Institutional Animal Ethics Committee (IAEC) of the Indian Institute of Science (IISc). A head post made of titanium was surgically implanted in both monkeys. It was then followed by a period of training for a visual fixation task. Post-training, a hybrid array of microelectrodes (Utah array), which consisted of 81 microelectrodes (9×9) and 9 ECoG (3×3) electrodes, was implanted in the primary visual cortex (V1) of the right cerebral hemisphere under anaesthesia. Microelectrodes (length = 1mm) had an interelectrode distance of 400 µm. Grid placement was approximately set at ≈ 15 mm lateral from midline in V1 and ≈ 15 mm rostral from the occipital ridge. The placement of reference wires was over the dura near the recording location, or they were wrapped around titanium screws on the surface of the skull near the craniotomy.

To get LFP data, the raw data obtained was band-pass filtered between 0.3 Hz (Butterworth filter, first order, analog) and 500 Hz (Butterworth filter, fourth order, digital), digitized at 16-bit resolution (sampled at 2kHz). Further, LFP signals were decimated by a factor of 8 (sampling frequency became 250Hz) using the MATLAB function ‘resample’ for this study.

### Behavioural task and Experimental design

The monkeys sat on a chair inside a Faraday cage enclosure to shield them from external electrical noise, and their heads were fixed by the head post while performing the visual fixation task. Monkeys were required to fixate their gaze within 2° of a small central dot (0.05° or 0.1° diameter) displayed on an LCD monitor screen (BenQ XL2411, LCD, 1280X720 pixels, 100 Hz refresh rate, gamma corrected). The monitor was placed 50 cm away from the eyes of the subjects. A single trial block consisted of an initial blank period of 1000 ms, followed by a display of a series of two to three stimuli in succession for 800 ms each, with an interstimulus interval of 700 ms. On successful fixation throughout the block, they were given a juice reward at the end.

Full-screen static gratings (covering ≈56° (width) and 33° (height) of the visual field) were the stimuli presented at full contrast, one of the eight orientations (0°, 22.5°, 45°, 67.5°, 90°, 112.5°,135° and 157.5°) and one of the five spatial frequencies (0.5, 1, 2, 4 and 8 cycles per degree). They were presented in pseudorandom order. The average number of trials for each pair of spatial frequency and orientation was 33 (28-36) for Monkey 1 and 42 (37-45) for Monkey 2. This dataset has been used in our previous studies (Dubey and Ray, 2020; Gautham and Ray, 2024; Prabhu and Ray, 2024). The receptive fields (RFs) of the neurons contributing to signals of microelectrodes were centered in the lower left quadrant of the visual field at an eccentricity of ∼3° to ∼4.5° in Monkey 1 and ∼1.4° to ∼1.75° in Monkey 2.

### Electrode selection

We limited our further analysis to electrodes, whose RF estimates were steady over the period of many days (SD less than 0.1°), similar to our earlier studies (Murty et al., 2018; Dubey and Ray, 2020). This resulted in 77 and 31 electrodes for Monkey 1 and Monkey 2, respectively.

## Data analysis

Our entire data analysis was performed using custom codes written in MATLAB R2024b (MathWorks, RRID: SCR_001622). For the construction of the time-frequency power spectrogram, we used a moving window of length 250 ms and a step size of 25 ms, leading to a frequency resolution of 4 Hz. For the power spectral densities (PSDs), the baseline period was defined as-500 to 0 ms of stimulus onset, and the stimulus period was considered to be 250 to 750 ms for each stimulus presentation to avoid stimulus-onset related transients in the first 250 ms. PSDs were computed for the stimulus and baseline periods using the multitaper method (single taper) using the Chronux toolbox (Mitra and Bokil, 2008) (http://chronux.org/, RRID: SCR_005547). The frequency resolution for estimated PSDs was 2 Hz.

The full-screen stimuli induced slow and fast gamma waves that depended on the orientation and spatial frequency of the stimuli. We restricted our entire analysis to stimuli with a spatial frequency of 1 CPD, as slow and fast gamma were simultaneously optimally induced across different stimulus orientations (Figure S1 B, D). Based on the average PSD traces, the slow gamma frequency range was considered to be between 20-32 Hz for Monkey 1 and 20-38 Hz for Monkey 2 (Figure S1 A, C). Similarly, the fast gamma frequency range was considered between 36-65 Hz for Monkey 1, and 42-65 Hz for Monkey 2. The power tuning curves of fast and slow gamma bands were constructed by averaging the power in these bands, as done in previous studies (Murty et al., 2018).

### Generation of synthetic LFP signal

For testing the performance of various burst detection methods, we generated synthetic LFP signals by injecting Gabor atoms (Gaussian-modulated sinusoidal waveforms, mimicking gamma bursts) of pre-determined fixed durations into the spontaneous LFP that was recorded when no stimulus was shown, similar to previous studies (Chandran Ks et al., 2018; Anand et al., 2025). In these previous studies, we had a condition where a stimulus of zero percent contrast (i.e., no stimulus) was presented, which was used to get long segments of ‘spontaneous’ LFP. Since such a condition was not present in the current dataset, we simulated long spontaneous LFP segments by interpolating the Fast Fourier Transform (FFT) of the baseline segment (500 ms duration). Specifically, to generate spontaneous LFP segments of longer durations, we interpolated the baseline LFP FFT amplitudes using the MATLAB function ‘interp1’, which were further normalized appropriately to keep the total energy fixed. Phase angles corresponding to the interpolated points were randomly generated. Next, we computed the inverse Fourier transform to generate the full-length spontaneous LFP segment in the time domain. Four times the standard deviation of an injected Gabor atom (covering around 95 % of the energy of the atom) was considered as the length of the burst. The center frequency of injected bursts was chosen from a uniform distribution between 20 and 65 Hz, as both the slow and fast gamma “bumps” were primarily localized in the defined range for both Monkeys. While injecting longer bursts (>250 ms), the number of bursts per trial was fixed to one. For bursts shorter than 250 ms, the number of bursts injected per trial was estimated from a Poisson distribution, with its expected value equal to the ratio of length of stimulus period (500 ms: 0.25 to 0.75 s) and length of bursts to be injected. The selection of the amplitude, time center, and other metrics of the injected synthetic burst was made using the same criteria as our previous study (Chandran Ks et al., 2018). Bursts of fixed durations ranging from 50 ms to 400 ms (with 50-ms step size) were injected. The average frequency spectrum of the synthetic LFP trials (separately for each burst length injected) was appropriately scaled with a constant to match the average PSD of the actual dataset, in the frequency range of 20-65 Hz.

### Burst detection methods Matching Pursuit algorithm (MP)

Matching pursuit (Mallat and Zhifeng Zhang, 1993; Durka et al., 2001; Chandran Ks et al., 2016, 2018) is an iterative signal decomposition-based algorithm. Adopting a greedy approach, it approximates a signal in the time domain as a linear combination of waveforms called atoms. Initially, an extensive and overcomplete dictionary of atoms (Gabor, Fourier and delta) is first made by scaling, shifting and modulating a single mother atom:

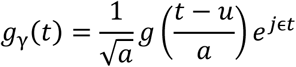

Here, 𝑔_γ_(𝑡) is an element (waveform or atom) of the dictionary (D), which we obtain by modulating the single mother atom 𝑔(𝑡) by γ = (𝑢, a, ɛ) that corresponds to the given values of parameters: translation (u), scale (𝑎), and modulation (ɛ). Normalization factor 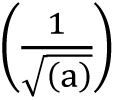 ensures that the energy of the atom is unity.

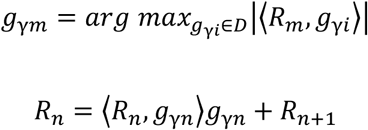

Here, 𝑓(𝑡) is the signal, while 𝑅_𝑚_ and 𝑔_γ𝑚_ are the residue and selected atom at the m^th^ iteration, respectively. At the initial iteration, the residue is set equal to the input signal itself (𝑅_0_ = 𝑓). This is followed by the computation of the inner product between the residue and all the atoms in the dictionary. The atom that has the largest inner product is selected. For subsequent iterations, the residue is set equal to the difference between the previous residue and the inner product between the chosen atom and the previous residue multiplied by the atom itself (approximation or projection). This procedure is repeated over a defined number of iterations. Upon completion of n iterations, the signal is decomposed as follows:

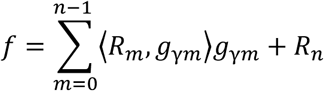

We used matching pursuit with a stochastic dictionary, in which the scale, shift, and modulation parameters are sampled uniformly. For our entire analysis, the dictionary size was 2.5 million, which was sufficient to sample the parametric space effectively (Chandran Ks et al., 2018; Anand et al., 2025). We used 120 iterations for the MP algorithm and achieved reasonable convergence (in terms of energy of the signal) with the given number of iterations. Prior to MP decomposition, the input LFP signal was band-pass filtered (using MATLAB function ‘filtfilt’) between 15 Hz and 80 Hz to decrease the required number of iterations and, therefore, to optimize the runtime of the algorithm. The scale parameter of a burst was used to compute its duration. Relevant bursts were selected according to their frequency center (within our frequency range of interest) and time center (within the stimulus period). Further, we filtered estimated bursts based on a power threshold criterion (relative to baseline) similar to one used in (Chandran Ks et al., 2018). This power threshold was specifically computed for the slow and fast gamma frequency ranges. We have previously shown that MP gives accurate results for a wide range of threshold values, unlike more traditional methods (see Figure 4 of (Chandran Ks et al., 2018)).

### Hilbert transform (HT)

For burst detection using the Hilbert transform (Lundqvist et al., 2016), the instantaneous power was considered as the square of the magnitude of the analytic signal, given as follows:

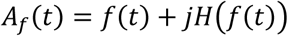

Here, 𝑓(𝑡)is defined as the band-pass filtered signal (in the frequency range of interest), using MATLAB function ‘filtfilt’ (Butterworth filter-order 4), and 𝐻 is the Hilbert transform. The instantaneous power was taken as the square of the instantaneous amplitude. Occurrences of gamma bursts (seed points) were located when the power exceeded three times the median power of the baseline signal (mBL). The start and end points of each burst were defined at the points where the power went below 1.5 times the mBL from that seed point, on either side of the seed point (similar metric as used in (Feingold et al., 2015) for beta burst detection).

### Orthogonal matching pursuit-Gabor expansion with atomic reassignment (OMP-GEAR)

OMP-GEAR is also an iterative algorithm, where it directly fits the residue (initially, the signal itself) to a new atom, using the magnitude of the inner product between the residue and the initial atom found by the canonical OMP method. It uses a unique parameter reassignment method for further refining the selected atom, thus not limiting the search space to only a finite-sized dictionary (for more details, see (Anand et al., 2025)). The dictionary size used for OMP-GEAR was 1.5 million, although smaller dictionaries also provide accurate results (Anand et al., 2025). Other specifics, such as the number of iterations (120), threshold criterion, and broad frequency range (15 Hz and 80 Hz) of initial band-pass filtering, were the same as those used for the MP algorithm. Rare occurrences of spurious bursts longer than 800 ms (stimulus duration) were excluded from the analysis across all methods.

### Analysis of burst dynamics and Statistical tests

For comparison of slow and fast gamma burst durations, we considered only those electrodes (for a given stimulus condition) for which more than 20 bursts were detected for both rhythms. Most of the electrodes that were previously selected based on receptive field estimation fulfilled the above criteria for both monkeys across all the methods.

The burst length distributions of slow and fast gamma were compared with a two–sample Kolmogorov-Smirnov (KS) test. The median durations and mean onset times (both electrode-wise) of slow and fast gamma bursts were analysed using the one-tailed two-sample Wilcoxon signed-rank test and paired t-test, respectively. Throughout our study, the standard error around the median was computed using the standard bootstrap method, with 1000 iterations.

The power matching for slow and fast gamma frequency ranges across the electrodes was performed using a histogram bin-height matching method (bin width: 0.5 dB (power)), as done in a previous study (Kumar and Ray, 2023). For each bin, we randomly selected electrodes (using MATLAB function ‘randsample’) from the rhythm having a longer bin height to make the bin height of both distributions equal. This resulted in two equal-sized groups of electrodes with similar fast and slow gamma power. Burst lengths of slow and fast gamma for the set of power-matched electrodes were compared using the one-tailed two-sample Wilcoxon rank-sum test across all methods.

### Connectivity estimation

We computed the phase locking value (PLV) or phase coherence (Lachaux et al., 1999), and the weighted phase lag index (WPLI) (Vinck et al., 2011) using the Fieldtrip Toolbox ((Oostenveld et al., 2011), RRID: SCR_004849). The PLV estimate between two signals from a pair of electrodes is defined as follows:

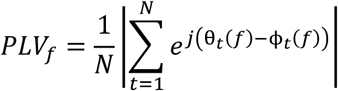

Here, 𝑁 denotes the total number of trials (in case of using a single taper), θ_𝑡_(𝑓) and ϕ_𝑡_(𝑓) are the phase angles of the estimated spectrum of the two signals, corresponding to 𝑓 (frequency) and 𝑡 (trial number). Further, we employed the WPLI metric in order to infer about the spatial scales of origin of the rhythm, as it has been shown to be a robust indicator of synchrony against noise and volume conduction (Vinck et al., 2011), and is computed as follows:

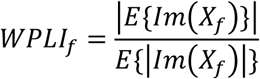

Here, 𝐼𝑚(𝑋_𝑓_) represents the imaginary component of the cross-spectrum between the two signals corresponding to f (frequency), and 𝐸 denotes the expected value or mean across trials. These connectivity measures were estimated for each pair of electrodes in the dataset. These pairs were further classified according to interelectrode distances. The baseline and stimulus periods were taken the same as above, resulting in a frequency resolution of 2 Hz. The number of tapers used for PLV and WPLI computation was 1 and 3, respectively.

### Model implementation details

We used a model implemented recently in our previous studies (Krishnakumaran and Ray, 2023; Krishnakumaran et al., 2025). It is an extension of the Jadi and Sejnowski (JS) model (Jadi and Sejnowski, 2014), which is a basic firing rate model consisting of an inhibitory and an excitatory population of neurons with a sigmoidal activation function, which, for a proper choice of inputs, operates as an Inhibition Stabilized Network (ISN). The model was able to generate gamma oscillation that replicates the decrease in peak frequency and increase in power with increasing size of the stimulus, as reported by previous studies in V1 (Gieselmann and Thiele, 2008), as well as an increase in the peak frequency with contrast (Ray and Maunsell, 2010). The population firing rates 𝑟_𝐸_ and 𝑟_𝐼_ of excitatory and inhibitory, respectively, evolve according to the following differential equations in the model:

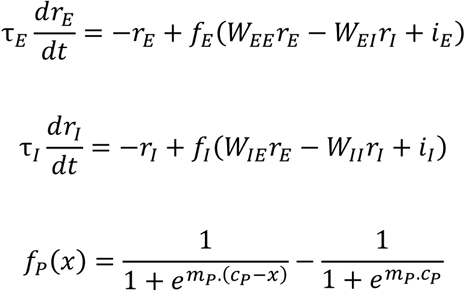

Here, the subscript P refers to the specific population, which could be either E (excitatory) or I (inhibitory). The original JS model produced sustained continuous limit cycles of gamma oscillations, as it employed constant input drives. Recruiting noisy input drives (specifically, Ornstein-Uhlenbeck (OU) noise) in the JS model has been shown to generate bursty gamma rhythm, with burst statistics similar to those observed in vivo (Krishnakumaran and Ray, 2023). The OU noise formulation of input drives 𝑖_𝐸_ and 𝑖_𝐼_ is given as follows:

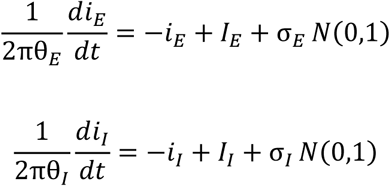

Here, 𝐼_𝐸_ and 𝐼_𝐼_ refers to the steady state mean values of 𝑖_𝐸_ and 𝑖_𝐼_ respectively, while N(0,1) denotes that at each time point, a value is generated from the standard Gaussian distribution. This formulation is essentially equivalent to additive white noise passed through a first-order low-pass filter, which has a cutoff frequency at θ_𝑃_ corresponding to the population P.

We independently generated slow and fast gamma by suitably varying τ_𝐼_, that is, the firing rate time constant of inhibitory neuronal populations, which determines the center frequency of the generated oscillation. For slow gamma, τ_𝐼_ was taken equal to τ_𝑆𝑂𝑀_ ≈ 15.4 ms, while for fast gamma, τ_𝐼_ was set as τ_𝑃𝑉_ ≈ 7.7 ms, where SOM and PV refer to somatostatin and parvalbumin-positive interneurons, respectively, which have been shown to play a role in specifically generating the two rhythms. These values of the firing rate time constant of the specific interneurons are similar to those used in previous studies (Garcia Del Molino et al., 2017; Krishnakumaran and Ray, 2023). Furthermore, 𝑊_𝐼𝐼_ for slow gamma simulation was set to zero, as there is generally no recurrent connection among SOM interneurons in V1 (Pfeffer et al., 2013; Garcia Del Molino et al., 2017; Bos et al., 2025). The remaining parameters were kept the same across the two simulations and were the same as used in our previous study (Tables 1 and 2, (Krishnakumaran and Ray, 2023)).

**Table 1:** Parameter values used in the model for generating fast gamma.

|  |  |
| --- | --- |
| $W_{EE}$ | 16 |
| $W_{EI}$ | 26 |
| $W_{IE}$ | 20 |
| $W_{II}$ | 1 |
| $m_E$ | 1 |
| $m_I$ | 1 |
| $c_E$ | 7.5 |
| $c_I$ | 18 |
| $\tau_E$ (ms) | 15.4 |
| $\tau_I = \tau_{PV}$ (ms) | 7.7 |

**Table 2:** Parameter values used in the model for generating slow gamma.

|  |  |
| --- | --- |
| $W_{EE}$ | 16 |
| $W_{EI}$ | 26 |
| $W_{IE}$ | 20 |
| $W_{II}$ | 0 |
| $m_E$ | 1 |
| $m_I$ | 1 |
| $c_E$ | 7.5 |
| $c_I$ | 18 |
| $\tau_E$ (ms) | 15.4 |
| $\tau_I = \tau_{SOM}$ (ms) | 15.4 |

We simulated the model with 128 distinct parameter combinations, obtained by varying the input drive (𝐼_𝐸_ = 𝐼_𝐼_ ∈ {7, 8, 9, 10, 11, 12}) and independently varying 𝜃_𝑃_ (𝜃_𝐸_, 𝜃_𝐼_ ∈ {1, 4, 16, 64}). As in our previous study, the variance parameter of the noise (𝜎 ^2^) was set equal to the steady-state mean input 𝐼_𝑃_. The range of parameters selected was similar to that in previous studies (Jadi and Sejnowski, 2014; Krishnakumaran and Ray, 2023), and to ensure that the slow and fast gamma were robustly generated within physiologically observed frequency ranges in V1.

Each parametric combination was simulated for 50 iterations for both slow and fast gamma, using the forward Euler method (time step size: 2×10^-5^). Each iteration had a timespan from - 2.047 to 2.048 s, where the input drives and variances were set to zero from-2.047 to 0 s, then instantaneously set to the desired non-zero value from 0 to 1.5 s, following which they were again set to zero. This is equivalent to a stimulus presentation duration of 1.5 s. We considered −𝑟_𝐸_ − 𝑟_𝐼_ as the proxy for LFP, similar to our previous studies (Krishnakumaran et al., 2022; Krishnakumaran and Ray, 2023). Further, it was passed through a fourth-order low pass Butterworth filter (cutoff frequency at 200 Hz) and downsampled by a factor of 200, resulting in a sampling frequency of 250 Hz (same as that in our actual dataset). Time-frequency spectra analysis and PSD estimation of the simulated model output were performed similarly to that described above for the actual LFP. Simulated slow gamma LFP proxy was band-pass filtered (using MATLAB function ‘filtfilt’) in the frequency range between 10 and 50 Hz, followed by MP analysis using 50 iterations. Fast gamma LFP proxy was filtered between 30 Hz and 90 Hz, followed by MP analysis using 100 iterations. The number of iterations was chosen to ensure that the MP algorithm achieves optimal convergence in decomposing the energy of the signal (band-pass filtered in the frequency range of interest).

Stimulus time period was defined as 0.25 to 1.25 s (to avoid stimulus-onset related transients). The slow gamma frequency range was considered to be between 20-40 Hz for Monkey 1, and the fast gamma frequency range was considered between 40-80 Hz. As mentioned above, relevant bursts were selected according to their frequency center (within our frequency range of interest), time center (within the stimulus period), and a power-based threshold. Any spurious bursts much longer than the stimulus duration were excluded from the analysis. For the model, due to the absence of a non-zero baseline power, the normalized power of the specific gamma band was estimated by dividing the maximum power (peak) in the frequency band by the average power of both ends of the frequency band (based on PSD), similar to a recent modelling study (Han et al., 2021a). Subsequently, power matching and other statistical analyses were performed across different parametric conditions in the same way as described above for the actual dataset.

## Code availability

The codes and data used for our entire analysis are available on the following GitHub repository: https://github.com/rvithebest/Slow_Fast_Gamma_Bursts.git

## Results

### Generation of Slow and Fast gamma rhythms by Large Visual Stimuli

We analysed LFP signals recorded from area V1 of two female monkeys while they viewed full-screen sinusoidal gratings of varying orientations and spatial frequencies. As shown previously, these full-field stimuli induced two distinct gamma rhythms, with distinct tuning preferences for both orientation (Figure S1) and spatial frequencies (Murty et al., 2018; Dubey and Ray, 2020). Figure 1A shows the average power spectral density (PSD; averaged over 280 stimulus repeats combined across all eight stimulus orientations) for an example electrode of Monkey 1 for which both slow and fast gamma rhythms were prominently induced, as evident by two distinct ‘bumps’ in the PSD (Figure 1A; magenta trace).

**Figure 1:**
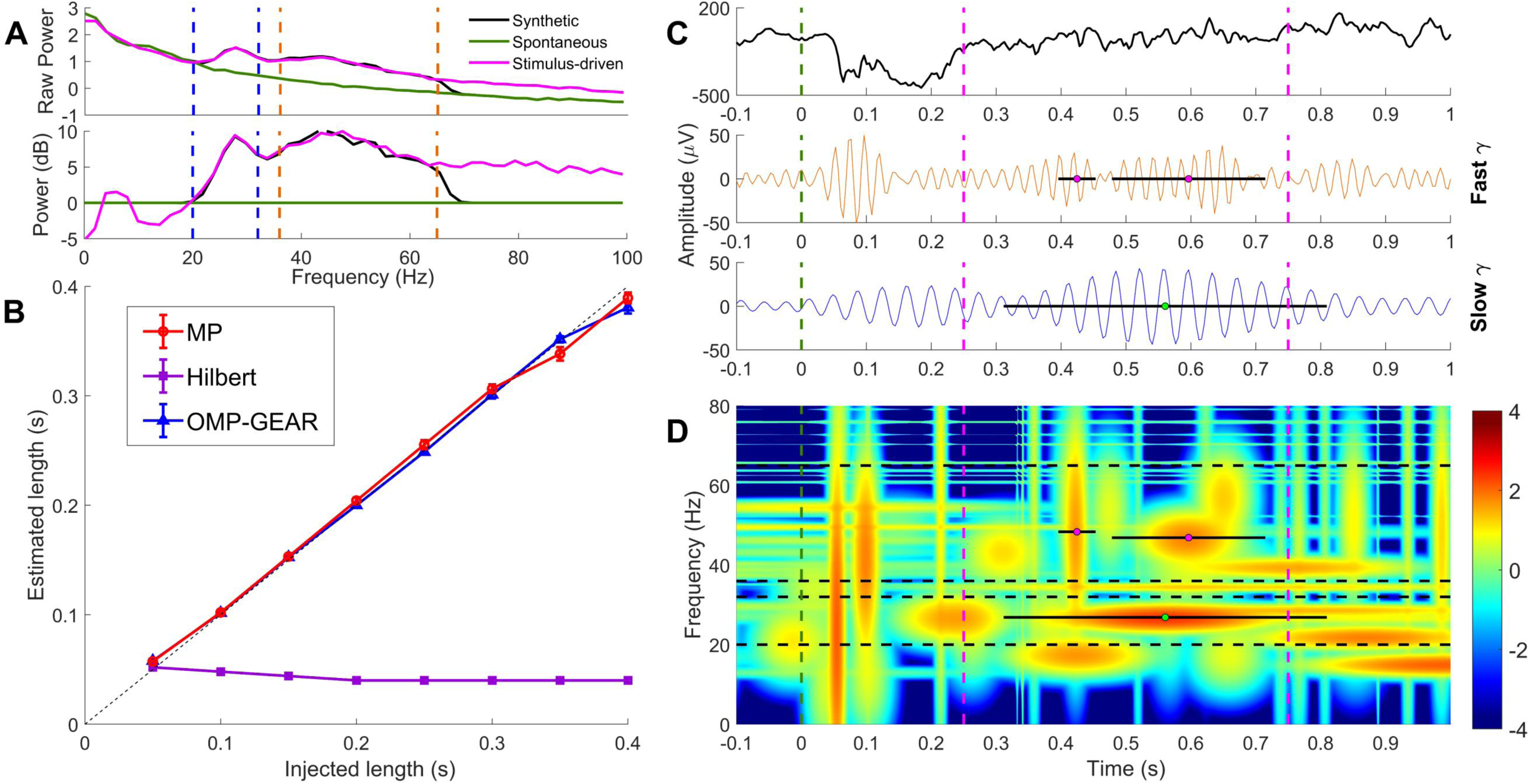
G**a**mma **burst estimation. (A)** Upper panel: Average power-spectral densities (PSD; log(µV^2^/Hz)) of three signals: Stimulus-driven (magenta trace), Spontaneous (green trace), and Synthetic LFP (black trace). Stimulus-driven signal corresponds to real LFP recordings from an example electrode in Monkey 1, for which full-screen, full-contrast stimuli were shown for 280 trials (combined across all orientations). PSD was computed between 250 to 750 ms after stimulus onset. The PSD for spontaneous LFP was obtained between-500 to 0 ms of stimulus onset. Synthetic LFP refers to the signal constructed by injecting bursts of fixed duration into each trial of spontaneous LFP (see Methods for details). The blue and orange dotted lines represent the frequency ranges of slow (20-32 Hz) and fast gamma (36-65 Hz), respectively for Monkey 1. Lower panel shows the change in power between stimulus and baseline (converted to ‘dB’ scale). **(B)** Performance of the three methods: MP (red), OMP-GEAR (blue), and HT (purple) on the synthetic LFP. The known burst lengths (50 ms to 400 ms, with 50 ms step size injected for 280 trials each) are plotted versus the median burst length along with the standard error of the median (SEM), as estimated by the three methods. The black dotted line denotes the ‘y=x’ line. **(C)** The top panel represents the raw LFP trace (black) of a single trial, from the same example electrode. The middle and bottom panels represent the same signal band-pass filtered in fast (orange) and slow gamma (blue) ranges, respectively. The black solid lines represent the burst durations detected by MP for this single trial (colored dot is the time center of the burst). **(D)** The time-frequency spectrum of the same single trial estimated by MP. Again, the black lines represent the burst durations detected by MP for this single trial. Vertical magenta dotted lines represent the stimulus period (0.25 to 0.75 s), while the vertical green dotted line indicates the stimulus onset (0 s). Horizontal black dotted lines represent the frequency ranges of slow (20-32 Hz) and fast gamma (36-65 Hz) for Monkey 1. Colorbar indicates the estimated power on a logarithmic scale.

### Robust burst length estimation by MP and OMP-GEAR

We first evaluated the performance of three methods (HT, MP and OMP-GEAR) for burst detection on the synthetic LFP signal, constructed by injecting Gabor atoms into spontaneous LFP (see methods for details). We injected Gabor atoms of pre-determined fixed durations (0.05 to 0.4 ms, with 0.05 ms step size), with center frequencies randomly selected within the broad frequency range of 20-65 Hz, thus mimicking either the physiological slow or fast gamma rhythms. These bursts were then suitably scaled to ensure that the average PSD of the synthetic LFP signal matched that of the real data in the frequency range of our interest (Figure 1A). More details about synthetic LFP generation can be found in our previous studies (Chandran Ks et al., 2018; Anand et al., 2025). Consistent with previous studies, MP and OMP-GEAR could robustly detect the injected bursts of varying durations, while the HT failed to efficiently compute burst lengths, especially for longer durations of bursts injected (Figure 1B, root mean square error (RMSE) value of fitting: 0.0071, 0.0076, 0.2171 for MP, OMP-GEAR and HT, respectively).

We proceeded with the MP algorithm to estimate burst durations in our dataset. Figure 1C shows the raw LFP trace from a single trial (black trace) and its bandpass filtered versions in the slow (blue trace) and fast (orange) gamma frequency range. The corresponding time-frequency spectrum is shown in Figure 1D. The gamma bursts identified by the MP algorithm in the slow and fast gamma ranges are shown as black line segments in Figure 1C and 1D. In this trial, we observed two short bursts of fast gamma and a single long slow-gamma burst.

### Longer burst lengths are observed for Slow gamma rhythm

Since both slow and fast gamma power depended on stimulus orientation (Figure S1), we first compared the burst durations for an orientation of 157.5°, for which power was comparable in the two gamma bands for Monkey 1 (for Monkey 2, slow gamma was weaker than fast at all orientations). The histogram of burst lengths combined across all electrodes for both the rhythms (Figure 2A, E) showed a characteristic peak (mode) at short burst durations, along with a right tail, consistent with previous studies where a single gamma rhythm was analyzed (Chandran Ks et al., 2018; Krishnakumaran and Ray, 2023).

**Figure 2:**
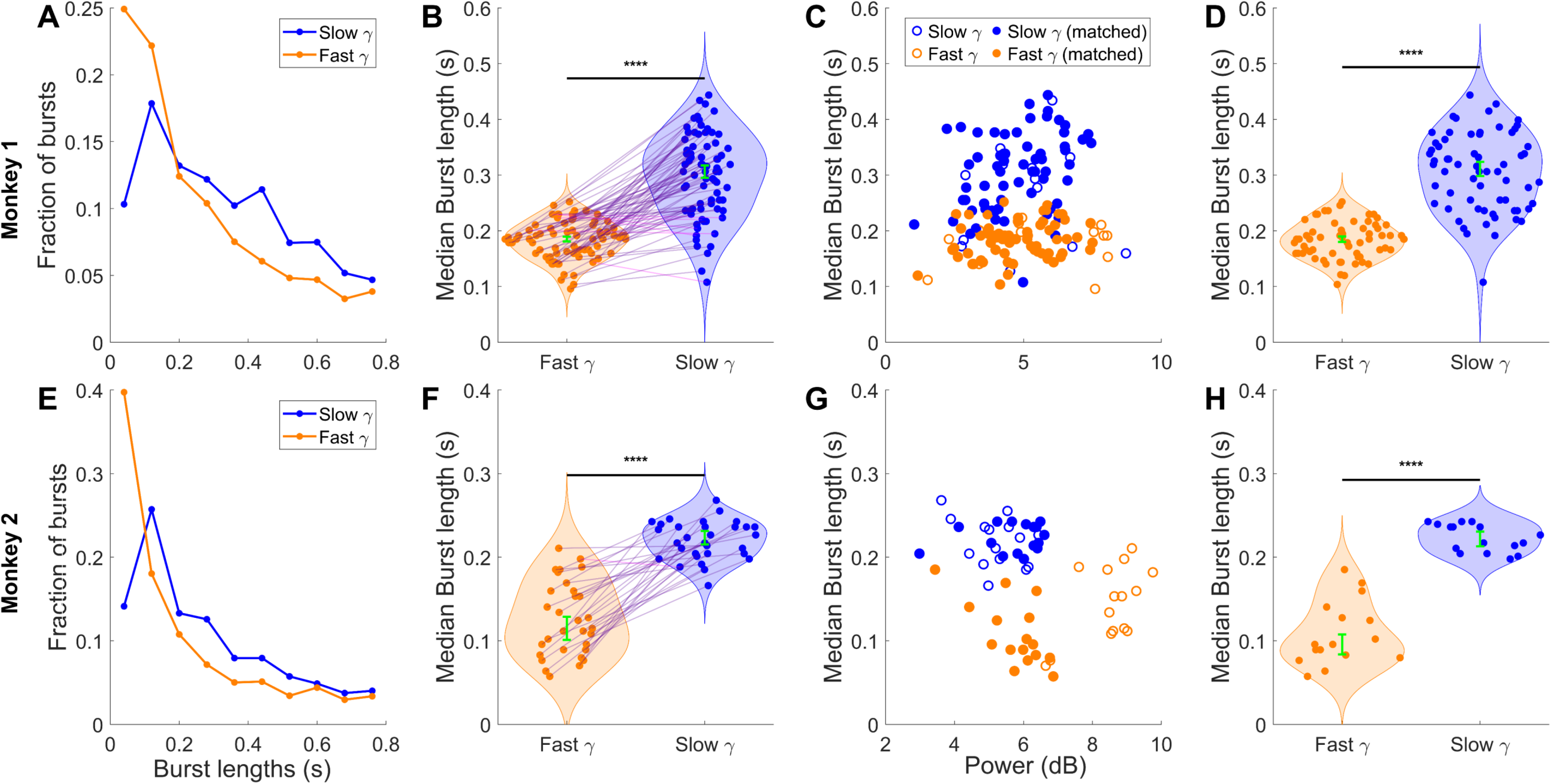
Comparison of burst durations of slow and fast gamma estimated by MP. (A, **E)** Histogram representing the distribution of burst durations of slow (blue) and fast (orange) gamma rhythm, combined across all electrodes of Monkey 1 and Monkey 2, when a stimulus with orientation of 157.5° and spatial frequency 1 CPD was shown. **(B, F)** Median burst lengths (electrode-wise) of slow and fast gamma (N=77 for Monkey 1 and N=31 for Monkey 2, where N denotes the number of electrodes considered for analysis). The green error bar represents the overall median along with the standard error of the median. **(C, G)** Median burst lengths of both rhythms plotted versus normalized power (dB) for all the sites of Monkey 1 and Monkey 2. Filled circles represent the set of randomly selected power-matched electrodes having similar power in slow and fast gamma. **(D, H)** Median burst lengths of slow and fast gamma for the power matched electrodes (N=64 for Monkey 1 and N=16 for Monkey 2 after power matching). The green error bar represents the overall median along with the standard error of the median.

Notably, there was a more pronounced heavy right-tail or right-shifted peak observed in the case of slow gamma, indicating that the proportion of longer burst durations observed was greater in the case of slow gamma as compared to fast gamma. The burst length distributions of the two rhythms were significantly different (KS-test; p-value equals 9.81 x 10^-80^ and 1.39 x 10^-109^ for monkey 1 and monkey 2, respectively). Further, the median burst durations for slow gamma rhythm exhibited significantly longer burst lengths than fast gamma for both monkeys (Figure 2B, F: Wilcoxon signed-rank test; N=77, p= 4.88 x 10^-14^, W-stat= 2968, Z-value= 7.44 for monkey 1 and N=31, p= 7.42 x 10^-7^, W-stat= 494, Z-value= 4.81 for monkey 2).

In our previous studies, we have observed a weak positive correlation between gamma burst durations and power in some cases, which was also observed for slow gamma in Monkey 1 (Spearman, r=0.28, p=1.35 x 10^-2^; Figure 2C; for other conditions, the correlations were not significant). To rule out the potential bias due to power differences between the rhythms, we considered a subset of electrodes having similar slow and fast gamma power (see Methods for details; the selected electrodes are indicated by solid circles in Figure 2C and 2G). Even after power matching, slow gamma rhythm exhibited significantly longer median burst durations as compared to fast gamma. (Figure 2D, H: Wilcoxon rank-sum test; N=64, p= 1.36 x 10^-17^, W-stat= 5903, Z-value= 8.46 for monkey 1 and N=16, p= 8.34 x 10^-7^, W-stat= 391.5, Z-value= 4.79 for monkey 2). The trend was consistent across all stimulus orientations, despite distinct power tuning curves exhibited by slow and fast gamma rhythms as mentioned above (Figure S2; Table S1).

### Analysis of time and frequency centers of bursts

Here, we analysed the temporal profiles of the slow and fast gamma bursts. Onset time for a given trial was defined as the time point at which the first (earliest) burst was detected/started. The estimated mean time-centers of the slow gamma bursts were significantly higher than those of fast gamma (Figure 3A, B, D, E; Two-sample paired t-test; N=77, p= 1.3881 x 10^-7^, t-stat= 5.640 for monkey 1 and N=31, p= 2.4196 x 10^-10^, t-stat= 9.0627 for monkey 2). Thus, we observed an increased latency for the onset of slow gamma bursts.

**Figure 3:**
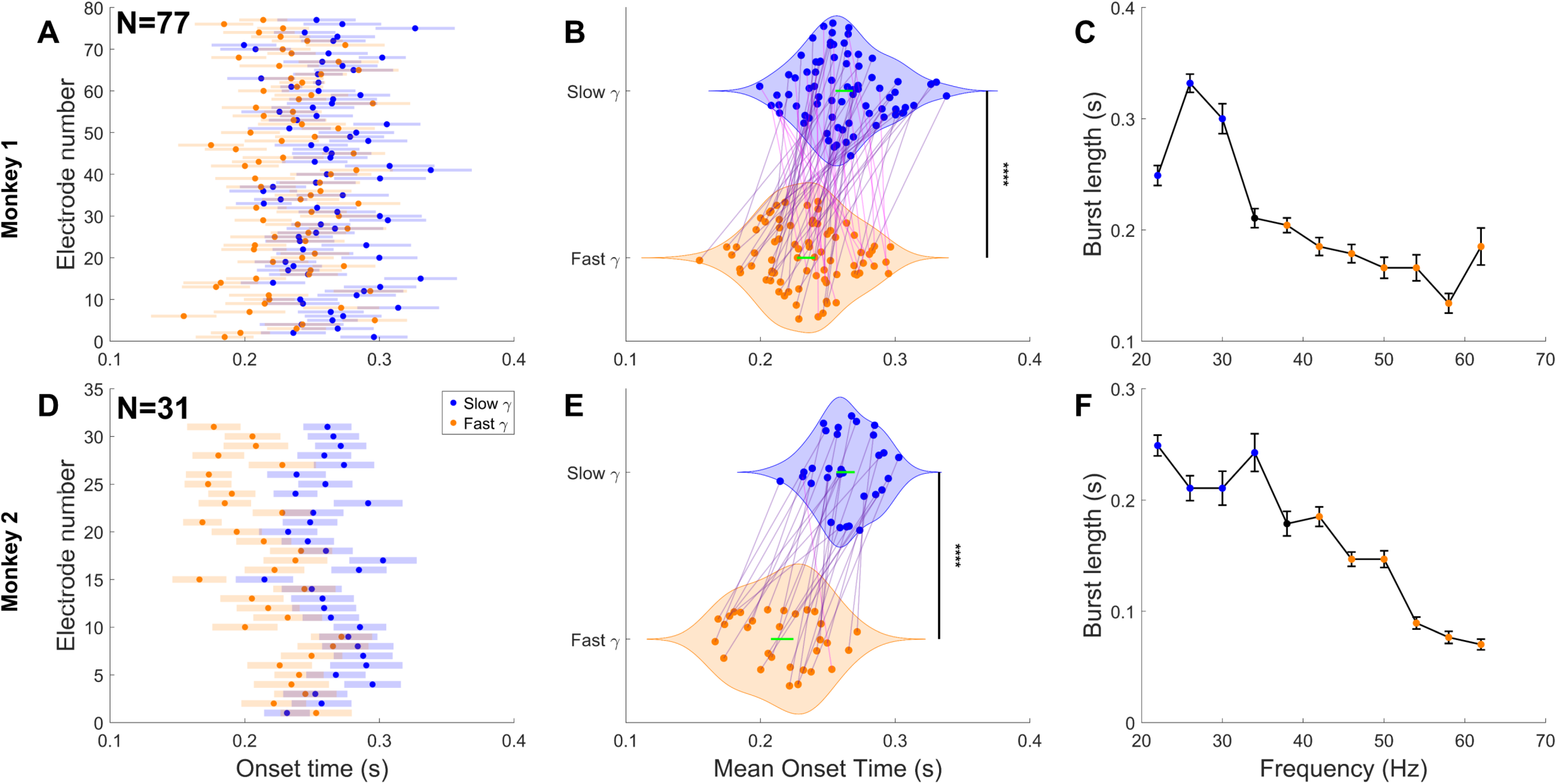
Analysis of the time and frequency centers of detected bursts. (A, **D)** Mean and standard error of the mean (SEM) of estimated onset times of the earliest observed burst in each trial for slow (blue) and fast (orange) gamma rhythm of Monkey 1 (N=77) and Monkey 2 (N=31) for stimulus orientation of 157.5°. **(B, E)** Mean onset times (electrode-wise) of slow and fast gamma bursts (N=77 for Monkey 1 and N=31 for Monkey 2). The green error bar represents the overall mean along with the SEM. **(C, F)** Estimated burst lengths are plotted versus their frequency center of the bursts (bin size equals 4 Hz). Error bars represent the median and standard error of the median of detected burst lengths combined across all sites. Frequency bin centers in slow and fast gamma frequency ranges are denoted in blue and orange centers, respectively.

We also tested the burst durations as a function of the center frequency of the burst. Bursts were consistently longer in the range corresponding to slow gamma rhythm, as compared to fast gamma. There was no major inflation or irregularity observed in burst lengths at the power line frequency (50 Hz), as our recordings were conducted in a heavily shielded Faraday cage, which aided in attenuation of line noise (Figure 3C, F). Similar results were observed for other methods as well, including HT and OMP-GEAR (Figure S3).

### Long-range synchrony mediated by slow gamma

We studied the functional connectivity of slow and fast gamma by computing the phase locking value (PLV) and weighted phase lag index (WPLI) profiles as a function of frequency, for trials corresponding to all the stimulus orientations, and electrode pairs were grouped according to the interelectrode distance (Figure 4A, C, E, G). While PLV is a measure of phase consistency across trials between two signals irrespective of the difference in phase angle, WPLI is typically used in EEG studies to reduce the effect of volume conduction because it measures the strength of non-synchronous (i.e., phase difference between signals that is different from zero) coupling (Vinck et al., 2011). Together, PLV and WPLI can be used to measure the strength of synchronous coupling between signals.

**Figure 4:**
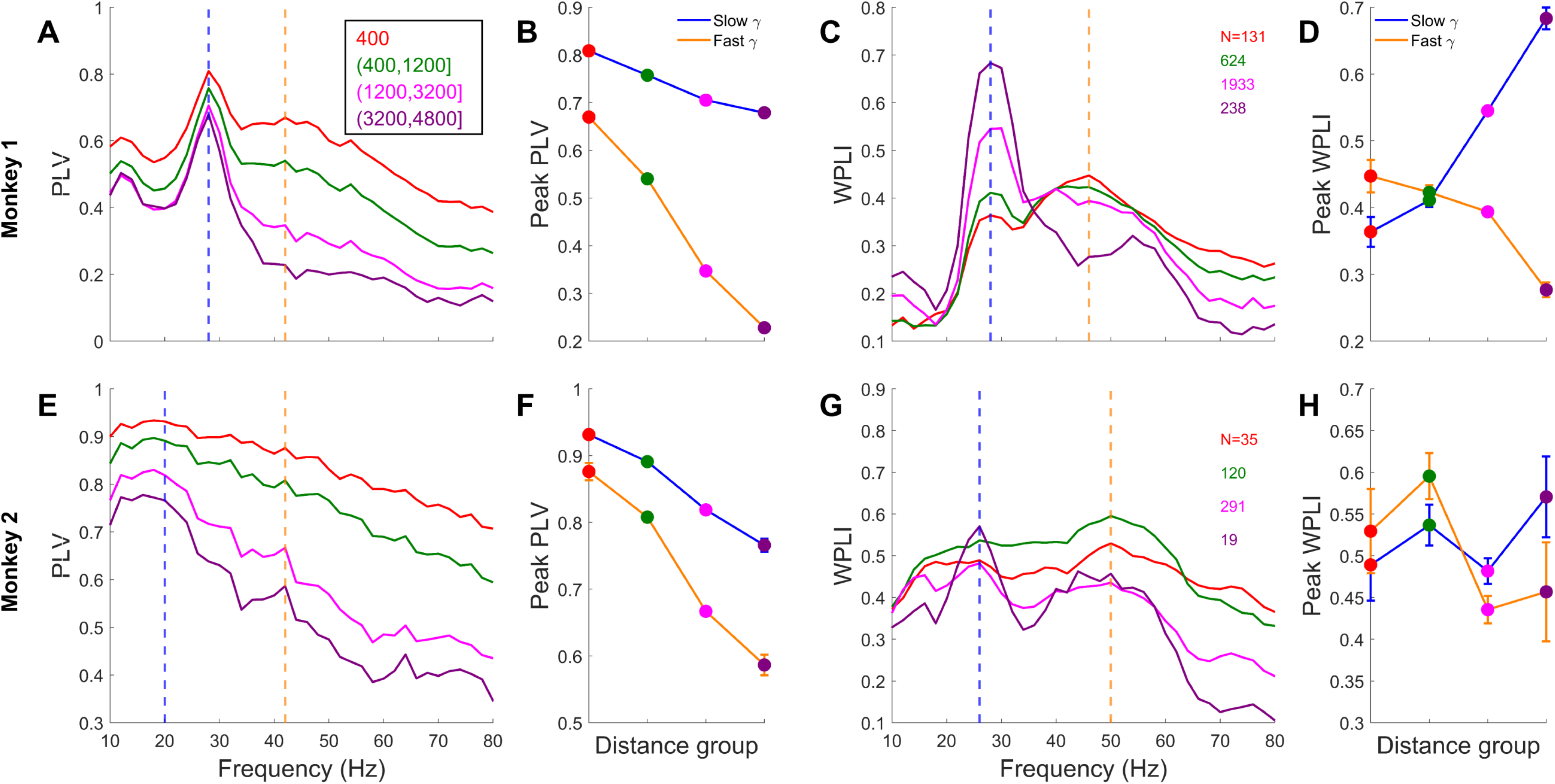
PLV and WPLI profiles of slow and fast gamma rhythm (A, **E)** Mean phase locking value (PLV) or phase coherence plotted against frequency, with electrode pairs grouped according to their interelectrode distance. Colored legends indicate the range of interelectrode distances considered for the corresponding group. The number of electrode pairs falling in the respective groups for Monkey 1 (N_Total_= ^77^C_2_= 2926) and Monkey 2 (N_Total_= ^31^C_2_= 465) are reported in **(C, G)** respectively (for all stimulus orientations combined). The blue dotted line indicates the frequency in the range of slow gamma corresponding to Peak PLV. The orange dotted line indicates the same in the fast gamma frequency range. **(C, G)** same as **(A, E),** for the weighted phase lag index (WPLI) method. **(B, F)** Average Peak-PLV values (taken at the indicated frequency) are plotted against the respective groups, with increasing interelectrode distances. The colored error bar represents the overall mean along with the standard error of the mean. **(D, H)** same as **(B, F),** for Peak WPLI estimates.

PLV remained high in the slow gamma range as the interelectrode distance increased, while it decayed more rapidly for fast gamma, as shown previously (Figure 4B, F; (Murty et al., 2018); Two-way ANOVA analysis with PLV estimates in the gamma bands and interelectrode distances as factors, resulted in a highly significant interaction; F_(3,5851)_ = 537.60, p= 1.52 x 10^-308^ for monkey 1 and F_(3,929)_ = 21.12, p= 3.08 x 10^-13^ for monkey 2). Interestingly, for slow gamma, WPLI values were much smaller than PLV at small inter-electrode distances and either increased (monkey 1) or remained the same (monkey 2) with increasing inter-electrode distances (Figure 4D, H; Two-way ANOVA analysis with WPLI estimates in the gamma bands and interelectrode distances as factors, resulted in a significant interaction; F_(3,5851)_ = 98.71, p= 2.53 x 10^-62^ for monkey 1 and F_(3,929)_ = 2.72, p= 0.04 for monkey 2). This suggests that slow-gamma coupling had a strong contribution from synchronous activity (zero phase-difference) at small inter-electrode distances. Unlike slow-gamma, fast-gamma WPLI also decreased with increasing inter-electrode distance in both monkeys, suggesting more local origins of fast gamma.

### JS Model with noisy inputs reproduces the differential burst dynamics

Finally, we tested whether these burst characteristics can be replicated in a stochastic JS model with inputs modulated by OU noise. To generate gamma oscillations at different frequencies, we varied the firing rate time-constant of inhibitory neuronal population to represent proxy SOM (𝜏_𝐼_ = 𝜏_𝑆𝑂𝑀_ =15.4 ms) and PV (𝜏_𝐼_ = 𝜏_𝑃𝑉_ =7.7 ms) interneurons similar to previous studies (Methods, (Garcia Del Molino et al., 2017)), which generated slow and fast gamma, respectively. Additionally, there were no recurrent connections among SOM interneurons (𝑊_𝐼𝐼_ = 0 for the slow gamma model; (Pfeffer et al., 2013)). Figure 5A, E summarizes the model schematics used for simulating the two rhythms, with parameter values as mentioned in Tables 1, 2, and Methods section. Average raw time-frequency spectrograms and power spectral densities (Figure 5B, C & G) demonstrate the generated slow and fast gamma rhythms in our modelling framework for a representative parametric combination. It verifies that, 𝜏_𝐼_ determines the center frequency of the simulated oscillation, as known from previous literature (Buzsáki and Wang, 2012).

**Figure 5:**
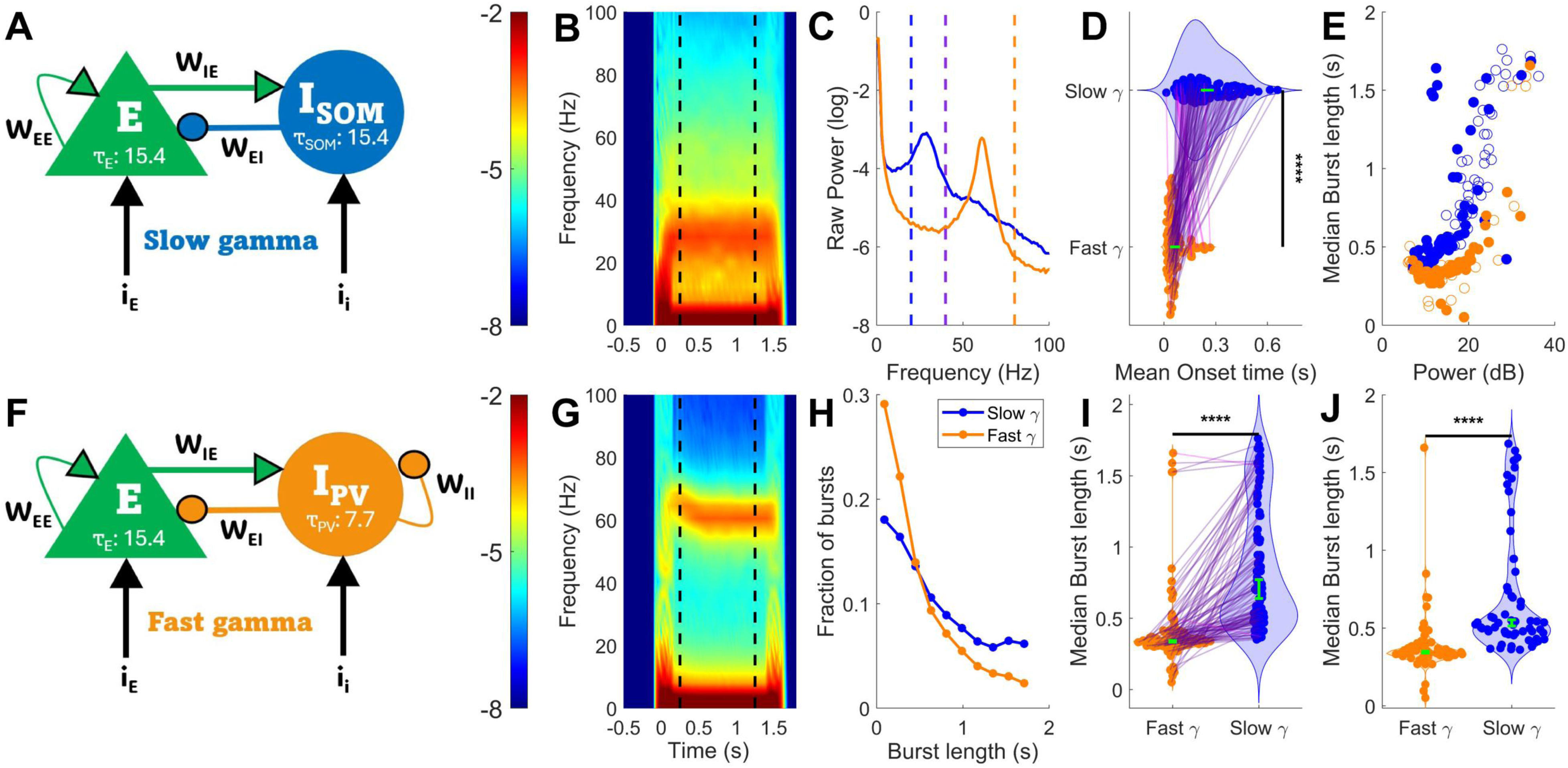
Analysis of simulated model outputs **(A, F)** The model schematics are used for generating slow and fast gamma rhythms, respectively. Full set of parameters are reported in Tables 1 and 2. E refers to excitatory population (pyramidal neurons), while SOM and PV, represent somatostatin and parvalbumin-positive interneuronal populations (inhibitory), respectively. **(B, G)** Average raw time-frequency plots (over 50 iterations) for simulated proxy LFP from the two models for given representative parametric combinations, confirming the generation of slow and fast gamma, respectively. Colorbar indicates the power (log-scale). Vertical black dotted lines represent the stimulus period (0.25 to 1.25 s). **(C)** Average raw power-spectral density plots (log-scale) from the two models (blue: slow gamma, orange: fast gamma) are shown for the same parameters given as **(B, G). (H)** Histogram representing the distribution of burst durations of slow (blue) and fast (orange) gamma rhythm estimated by MP across all parametric conditions**. (D)** Mean onset times (across distinct parametric combinations (N=105), with 50 iterations each) of slow and fast gamma bursts. The green error bar represents the overall mean along with the standard error of the mean. **(I)** Median burst lengths (across distinct parametric combinations (N=105), with 50 iterations each) of slow and fast gamma. The green error bar represents the overall median along with the standard error of the median. **(E)** Median burst lengths of both rhythms plotted versus normalized power (dB) for all the specified parametric combinations. Filled circles represent the set of randomly selected power-matched conditions, having similar power of slow and fast gamma. **(H)** Median burst lengths of slow and fast gamma are represented for the power-matched conditions (N=59). The green error bar represents the overall median along with the standard error of the median.

Subsequently, we performed similar MP analysis on the simulated proxy LFP. We used 128 different parametric combinations, spanning a wide range of mean input drives and OU noise parameters, in order to effectively vary the power and spectral variance of the generated rhythms (see Methods). Interestingly, simply changing the inhibitory time constant was sufficient to replicate other characteristics of the bursts. First, a more prominent heavy tail was observed for slow gamma as compared to fast gamma rhythm in the burst duration distributions of the two rhythms, which were significantly different (Figure 5H; KS-test; p-value less than 10^-308^, combined across all parametric combinations). Second, slow gamma exhibited longer median burst lengths and delayed onset than fast gamma in the modelling analysis across the different combinations. Bursts of both slow and fast gamma rhythm were detected in 105 parametric combinations (Figure 5D, I; Two-sampled paired t-test for mean onset time; N=105, p= 2.40 x 10^-21^, t-stat= 11.86; Wilcoxon signed-rank test for median burst length; N=105, p= 7.93 x 10^-19^, W-stat= 5530.5, Z-value= 8.78). As in real data, median burst lengths were higher for slow gamma even after performing power matching (The selected electrodes are indicated by solid circles in Figure 5E; Figure 5J: Wilcoxon rank-sum test; N=59, p= 8.20 x 10^-14^, W-stat= 4881, Z-value= 7.38).

Parameters of OU noise (𝜃_𝐸_, 𝜃_𝐼_) also influence the burst durations of the rhythm by dictating the temporal dynamics of the input in our framework (Krishnakumaran and Ray, 2023). We observed that the absolute median burst length of both slow gamma and fast gamma generally decreased with either increasing 𝜃_𝐸_ or 𝜃_𝐼_ (Figure S4), as shown previously (Krishnakumaran and Ray, 2023). Importantly, when compared at an identical set of parameters of noisy inputs, median duration of slow gamma remained longer than fast gamma.

## Discussion

We compared the burst statistics of stimulus-induced slow and fast gamma oscillations in the primary visual cortex of non-human primates using several burst detection algorithms. We observed longer bouts and increased latency of stimulus-induced slow gamma rhythm as compared to fast gamma. The two rhythms had differential PLV and WPLI profiles, with higher long-range synchronization in slow gamma range compared to fast gamma. Surprisingly, simply increasing the inhibitory firing rate time-constant in a WC model operating in an ISN range was sufficient to not only produce slower rhythms, but also increase their duration and onset latency. Thus, simply slowing down the rhythm also made it longer and delayed its onset.

### Distinct roles of slow and fast gamma and importance of burst statistics

The physiological gamma rhythm burst statistics were shown to be generated in a specific optimum parametric space based on a stochastic neural network, with variations in network connectivity parameters leading to pathological burst statistics (Powanwe and Longtin, 2019). A few other theoretical studies have emphasized the role of transient gamma bursts in inter-areal communication across the cortex (Lowet et al., 2017; Palmigiano et al., 2017). Therefore, the observed differential burst properties of slow and fast gamma suggest that the two rhythms have separate functional roles and distinct neural correlates. These two gamma oscillations have been shown to exhibit travelling waves in Macaque V1 (Gautham and Ray, 2024), which propagate in different uncorrelated directions. The longer lifetime of travelling waves produced by slow gamma, in comparison to fast gamma, is in accordance with our findings. We also observed distinct power-tuning curves for the two rhythms across stimulus orientations and spatial frequency, which is consistent with previous studies indicating enhanced information encoding in the cortex with the presence of two rhythms (Murty et al., 2018; Han et al., 2021b). We recently found that while fast gamma phase is coupled to power in frequencies above 150 Hz (which in turn is tightly coupled with multiunit firing around the microelectrode (Ray and Maunsell, 2011), slow gamma shows phase-amplitude coupling to frequencies between 80-150 Hz (Prabhu and Ray, 2024). These results are consistent with the hypothesis that fast gamma is important for local processing and strongly couples with spiking activity in a local network, while slow gamma has a more modulatory influence, potentially involved in locking of dendritic spikes (Prabhu and Ray, 2024).

The inferences from our estimated PLV and WPLI profiles also indicate that fast and slow gamma rhythms mediate processing at distinct spatial scales, i.e., local and global, respectively. The field-field coherence measurements have been shown to be substantially higher for slow gamma than for fast gamma in studies in both mice and macaque V1, even for widely separated electrodes (Veit et al., 2017; Murty et al., 2018). Slow gamma was shown to be generated by long-range horizontal connections in a modelling study (Han et al., 2021a). These observations, along with the above-mentioned experimental evidence regarding the biophysical origins of the rhythms, lead to a hypothesis for slow gamma serving as a global modulatory signal (Veit et al., 2023). Longer bouts of slow gamma lead to an increased number of oscillatory cycles per burst, which could potentially compensate for its lower frequency (Zheng et al., 2016) and engage large populations of neurons across the neocortex for extended durations. On the contrary, shorter transient bursts of fast gamma (typical gamma oscillation) could be optimum for local synchronisation and influencing spikes, as indicated by a modelling study (Saraf and Young, 2021).

### Implications regarding biophysical mechanisms

Experimental studies in mouse V1 employing optogenetics have tried to decipher the specific mechanisms behind the generation of these distinct gamma rhythms (Chen et al., 2017; Veit et al., 2017). The perisomatic inhibition by parvalbumin-positive (PV+) interneurons has been shown to contribute to the production of fast gamma. On the other hand, the generation of slow gamma rhythm is mediated by the dendrite-specific inhibition provided by somatostatin (SOM) interneurons. Dendrites-specific inhibition by SOM interneurons leads to considerably slower negative feedback (E-I network), as compared to that provided by direct perisomatic inhibition by PV+ neurons (which acts as a shunt, directly inhibiting spike production). Furthermore, a study in the mouse hippocampus has shown that the kinetics of inhibitory post-synaptic currents (IPSC) mediated by GABA_A_ receptors is slower (latency in opening and closing) in the dendrites, as compared to that in the soma of pyramidal cells (Huang et al., 2023). Similar variability in the properties of post-synaptic receptors along the somato-dendritic axis might be present in pyramidal cells of V1 as well. Therefore, these could be the plausible biophysical mechanisms behind the increased latency and longer durations of slow gamma bursts. However, other than the synaptic currents, active dendritic spikes (Ca^2+^ and NMDA), which exhibit slower and sustained kinetics, could also contribute to the extended bouts of slow gamma oscillation. Despite slow gamma rhythm exhibiting a considerably higher proportion of longer burst lengths, the mode of the distribution still lies around 100 ms. We suspect the heterogeneity in the distribution of slow gamma burst durations could be linked to the rich repertoire of active dendritic conductances, which in turn exhibit variability across space and plasticity with time (Sinha and Narayanan, 2022).

### Modelling studies

Recently, an in-silico study recruiting a morphologically realistic 3D model of a pyramidal neuron in V1 has demonstrated that the dendrite-targeting rhythmic inhibition (specifically at slow gamma frequencies) efficiently modulates the dendritic Ca^2+^ and NMDA spikes (dendritic Na^+^ spikes were unaffected; (Headley et al., 2024)). Examining the putative reciprocal influence of dendritic currents on the generation of slow gamma in a modelling framework could be insightful, although it would require intensive computational resources. An alternative strategy is to employ simplified phenomenological network models, as done in this study. We were able to replicate the differential burst dynamics of slow and fast gamma in a stochastic JS model framework. Bursts manifested as noise-interrupted limit cycles in our model, and thus primarily a larger value of firing-rate time constant (𝜏_𝐼_) defined for SOM as compared to PV inhibitory interneurons, lead to reduced susceptibility to noisy inputs, subsequently generating sustained longer bursts of slow gamma rhythm. Longer latency to the onset of the slow gamma bursts also followed trivially from the larger 𝜏_𝐼_, since it took longer for the system to move to the regime where it produced salient gamma oscillations when the time-constant was large. While the absolute value of median burst durations was influenced by the given parameters of the OU noise-mediated inputs (Krishnakumaran and Ray, 2023), the trend between the two rhythms remained consistent irrespective of parametric conditions. Thus, the 𝜏_𝐼_ values assigned to SOM and PV interneuron were similar to those used in a previous study (Garcia Del Molino et al., 2017) and determined the center frequency (Buzsáki and Wang, 2012) of the generated oscillation as well as the temporal evolution of its bursts in our model.

We kept the two inhibitory populations non-interacting, since our main goal was to test whether the burst characteristics could be explained by simply changing a time constant in the model. Previous studies have implemented network models with multiple inhibitory neuronal populations (two or three), such as a stabilized supralinear network regime in case of firing-rate models or employing spiking neural networks to propose specific functions of these inhibitory interneurons in mediating circuit dynamics (Ter Wal and Tiesinga, 2021; Veit et al., 2023; Holt et al., 2024; Bos et al., 2025). It could be possible to suitably modify the WC-type models to generate multiple gamma rhythms concurrently, manifesting as bursts and thereby validate our results in another framework. However, due to the asymmetric inhibition of PV by the SOM interneurons in V1 (Pfeffer et al., 2013) and incorporating the effects of modulatory inputs, competing slow and fast gamma rhythms could arise in such a three-population network models that interact with each other in non-trivial ways (Keeley et al., 2017). Such models could explain how multiple classes of interneurons could synergistically modify spiking activity in a network.

### Burst profiles as potential diagnostics

The causal influence of gamma rhythm on cognitive processes and well-being remains a matter of debate, as the existing evidence is mostly correlational. However, the potential role of these burst profiles as biomarkers for the diagnosis of cognitive deficits could not be ruled out. For example, burst properties of the beta rhythm have been shown to change significantly in subjects with Parkinson’s disease as compared to healthy controls (Vinding et al., 2020). Previous studies have extensively focused on the decline of trial-averaged power of gamma oscillations computed using conventional spectral analysis in subjects with neurological disorders (An et al., 2018; Murty et al., 2021), neglecting the bursty nature of gamma rhythm. Moreover, stimulus-induced slow gamma rhythm appears to be quite salient in Human EEG recordings, potentially due to its global nature (Murty et al., 2018). Therefore, analyzing the burst statistics of slow gamma rhythm in EEG or MEG recordings and examining whether they change with the onset of cognitive impairments could be an intriguing question for the future.

---

**Figure S1:**
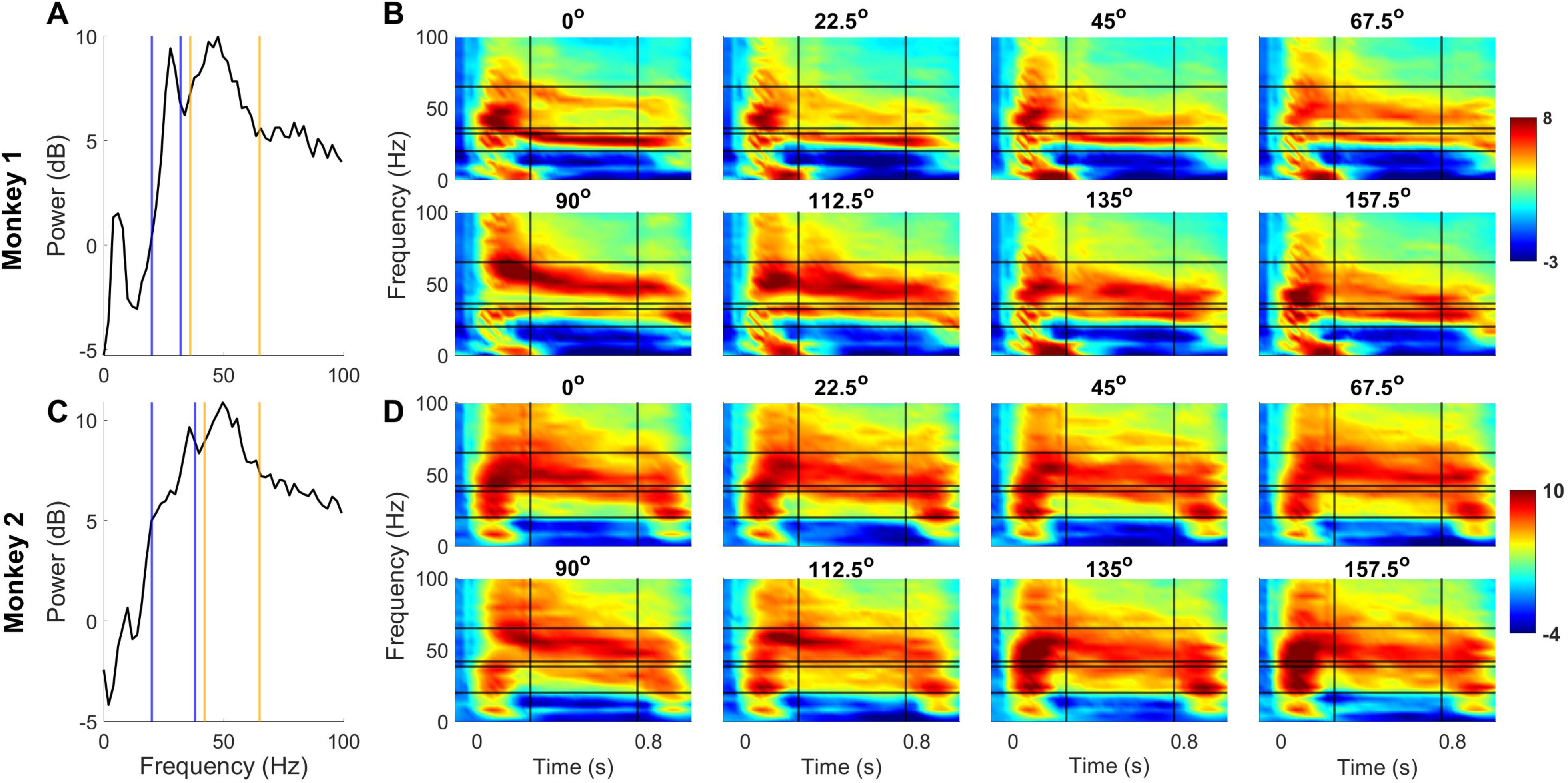
Representative average PSD traces and time-frequency spectrograms. **(A)** Average normalized power-spectral density (with respect to baseline, dB scale) of LFP recordings from an example electrode in Monkey 1, for 280 trials corresponding to stimuli of all orientations and spatial frequency of 1 CPD (Same as red trace of Figure 1B). **(C)** same as **(A)**, for Monkey 2. The blue and orange solid lines represent the considered frequency ranges of slow and fast gamma, respectively, for the corresponding monkeys. **(B)** Normalized time-frequency spectrogram (with respect to baseline, colorbar: Power (dB scale)), for each of the eight orientations averaged across all the electrodes for Monkey 1. **(D)** same as **(B)**, for Monkey 2.

**Figure S2:**
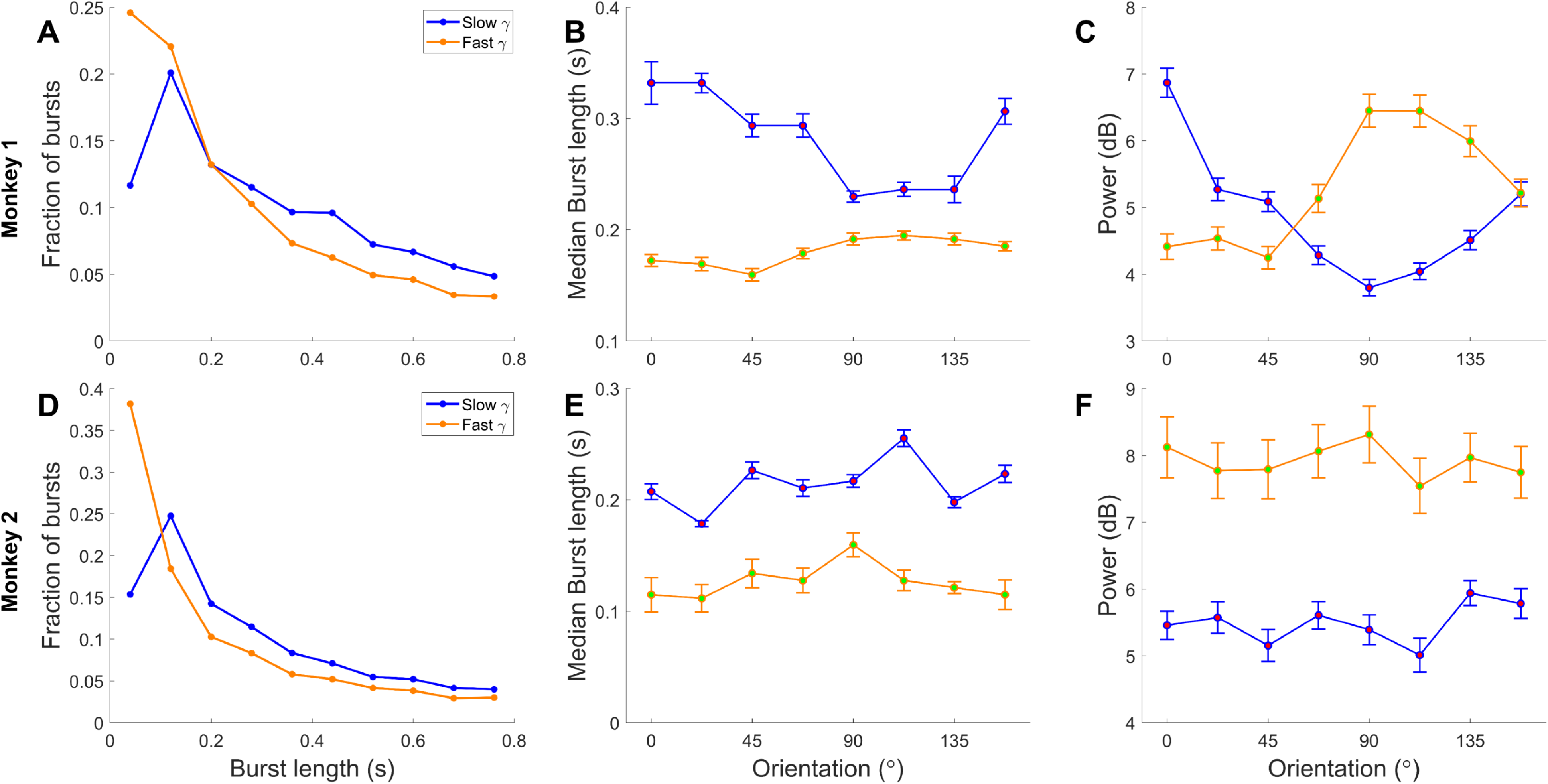
Burst durations and power tuning across all stimulus orientations. (A, **D)** Histogram representing the distribution of burst durations of slow (blue) and fast (orange) gamma rhythm, combined across all electrodes and orientations of Monkey 1 and Monkey 2. A significantly higher fraction of longer bursts is observed for the slow gamma as compared to fast gamma rhythm (KS-test; p-value less than 10^-308^). **(B, E)** Median and standard error of median of median burst lengths (electrode-wise) of slow and fast gamma for each orientation for Monkey 1 (N=77) and Monkey 2 (N=31). Median slow gamma burst durations were significantly longer than those of fast gamma at all orientations (Wilcoxon sign-rank; corresponding statistics are reported in Table S1). **(C, F)** Normalized power tuning curves (dB) across different stimulus orientations of Monkey 1 and Monkey 2. Error bars indicate the mean and SEM (standard error of the mean) of power values across electrodes.

**Figure S3:**
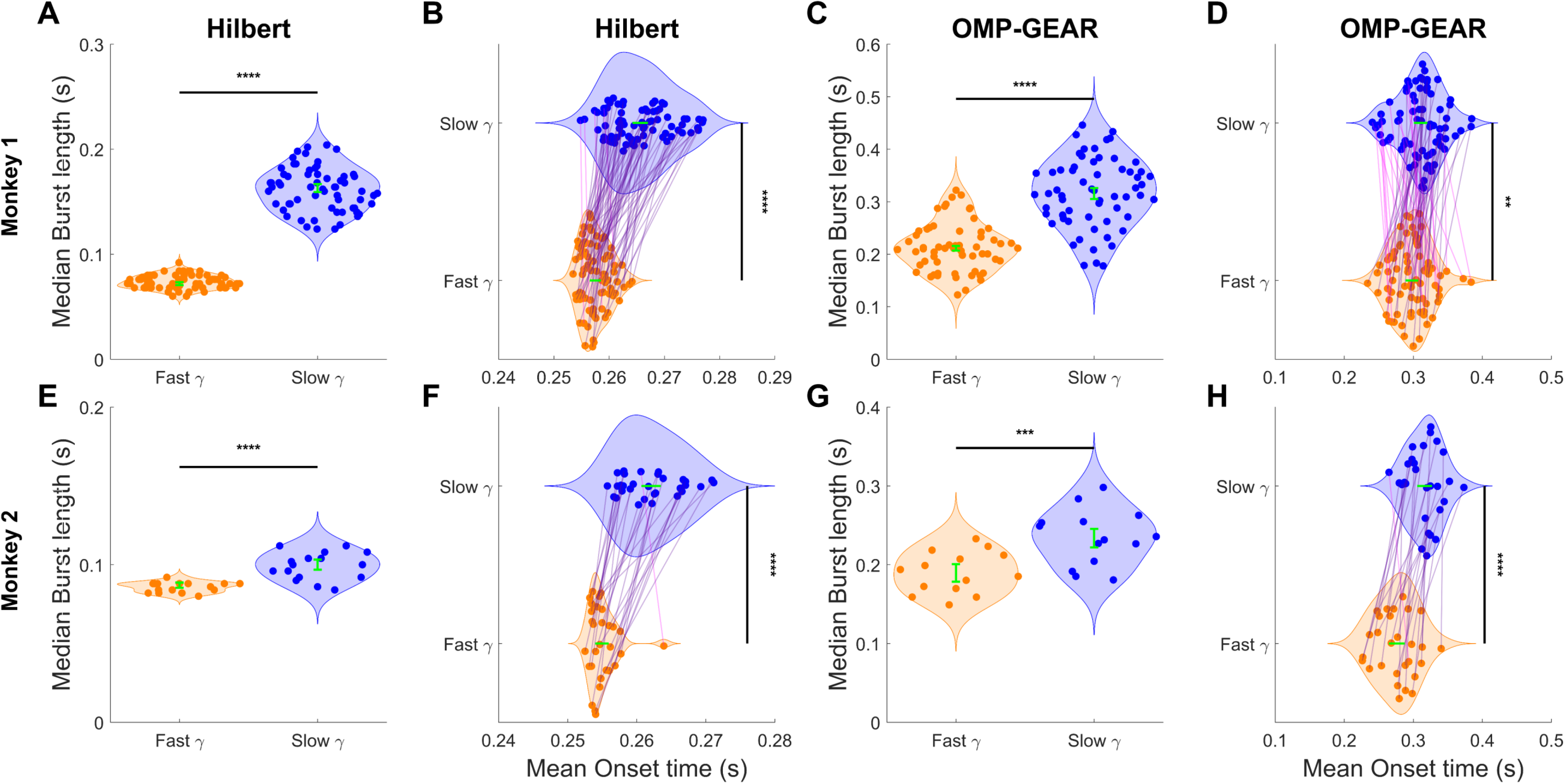
Results using HT and OMP-GEAR. **(A, E)** Median burst lengths of slow and fast gamma computed by HT for the power-matched electrodes (Wilcoxon rank-sum test; N=64, p= 8.22 x 10^-23^, W-stat= 6176, Z-value= 9.76 for monkey 1 and N=16, p= 3.09 x 10^-5^, W-stat= 370.5, Z-value= 4.01 for monkey 2; for stimulus orientation equal to 157.5°). The green error bar represents the overall median along with the standard error of the median. **(C, G)** same as **(A, E),** for the OMP-GEAR method (Wilcoxon rank-sum test; N=61, p= 3.04 x 10^-14^, W-stat= 5218, Z-value= 7.51 for monkey 1 and N=14, p= 9.63 x 10^-4^, W-stat= 271, Z-value= 3.10 for monkey 2). **(B, F)** Mean onset times (electrode-wise) of slow and fast gamma bursts computed by HT for all electrodes (Two-sampled paired t-test; N=77, p= 3.73 x 10^-21^, t-stat= 12.91 for monkey 1 and N=16, p= 4.51 x 10^-10^, t-stat= 8.76 for monkey 2). The green error bar represents the overall mean along with the standard error of the mean. **(D, H)** same as **(B, F),** for the OMP-GEAR method (Two-sampled paired t-test; N=74, p= 2.60 x 10^-3^, t-stat= 2.88 for monkey 1 and N=28, p= 7.04 x 10^-8^, t-stat= 7.05 for monkey 2).

**Figure S4:**
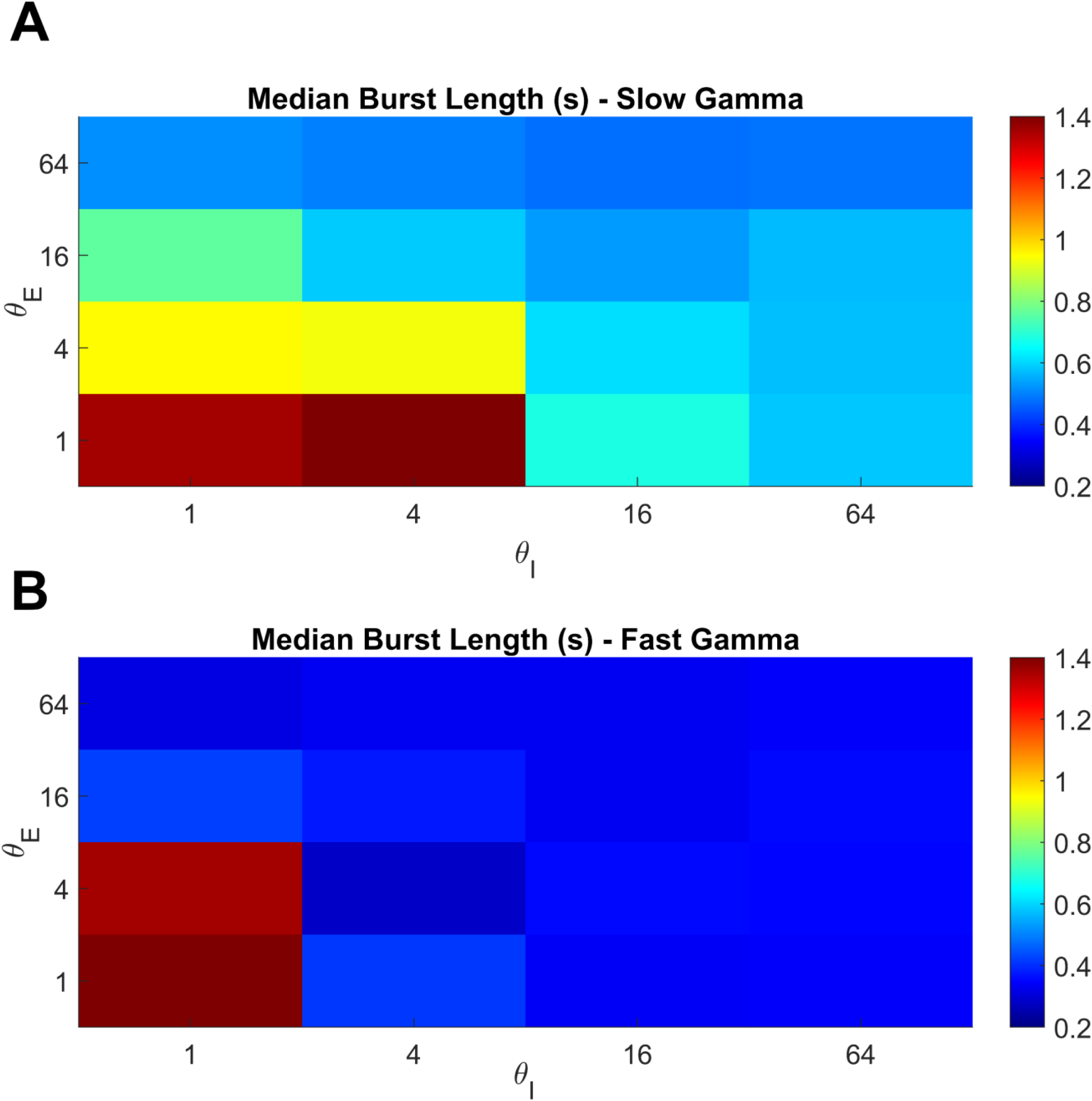
Median burst duration of slow and fast gamma rhythms as a function of 𝜽_𝑬_ **and** 𝜽_𝑰_ **(A)** Median burst duration of slow gamma in seconds (colorbar) as a function of 𝜃_𝐸_ and 𝜃_𝐼_ (combined across all values of steady state input current). **(B)** Same as **(A)**, but for fast gamma.

**Table S1:** Wilcoxon signed-rank test was performed for each of the orientations (median burst durations (electrode-wise)) as shown in. **Figure 3B, E**

| **Monkey-1 (N=77)** |  |  |  | **Monkey-2 (N=31)** |  |  |
| --- | --- | --- | --- | --- | --- | --- |
| **Orientation (°)** | **p-value** | **Z-value** | **W-stats** | **p-value** | **Z-value** | **W-stats** |
| **0** | $2.42 \times 10^{-14}$ | 7.54 | 2919 | $6.14 \times 10^{-7}$ | 4.85 | 496 |
| **22.5** | $1.39 \times 10^{-13}$ | 7.30 | 2940.5 | $1.42 \times 10^{-6}$ | 4.68 | 460.5 |
| **45** | $5.27 \times 10^{-14}$ | 7.43 | 2966 | $1.66 \times 10^{-6}$ | 4.65 | 459 |
| **67.5** | $2.68 \times 10^{-13}$ | 7.22 | 2923 | $1.66 \times 10^{-6}$ | 4.65 | 433 |
| **90** | $2.16 \times 10^{-8}$ | 5.48 | 2462.5 | $2.71 \times 10^{-6}$ | 4.55 | 480.5 |
| **112.5** | $3.84 \times 10^{-7}$ | 4.94 | 2418 | $8.67 \times 10^{-7}$ | 4.78 | 492.5 |
| **135** | $6.77 \times 10^{-11}$ | 6.42 | 2703.5 | $2.83 \times 10^{-6}$ | 4.54 | 480 |
| **157.5** | $4.88 \times 10^{-14}$ | 7.44 | 2968 | $7.42 \times 10^{-7}$ | 4.81 | 494 |

